# Training propulsion: Locomotor adaptation to accelerations of the trailing limb

**DOI:** 10.1101/582841

**Authors:** Andria J. Farrens, Rachel Marbaker, Maria Lilley, Fabrizio Sergi

## Abstract

Many stroke survivors suffer from hemiparesis, a condition that results in impaired walking ability. Walking ability is commonly assessed by walking speed, which is dependent on propulsive force both in healthy and stroke populations. Propulsive force is determined by two factors: ankle moment and the posture of the trailing limb during push-off. Recent work has used robotic assistance strategies to modulate propulsive force with some success. However, robotic strategies are limited by their high cost and the technical difficulty of fitting and operating robotic devices with stroke survivors in a clinical setting.

We present a new paradigm for goal-oriented gait training that utilizes a split belt treadmill to train both components of propulsive force generation, achieved by accelerating the treadmill belt of the trailing limb during push off. Belt accelerations require subjects to produce greater propulsive force to maintain their position on the treadmill and increases trailing limb angle through increased velocity of the accelerated limb.

We hypothesized that accelerations would cause locomotor adaptation that would result in measurable after effects in the form of increased propulsive force generation. We tested our protocol on healthy subjects at two levels of belt accelerations. Our results show that 79% of subjects significantly increased propulsive force generation, and that larger accelerations translated to larger, more persistent behavioral gains.

## I. INTRODUCTION

Eight out of ten stroke survivors suffer from hemiparesis, a condition that causes unilateral muscle weakness that results in gait asymmetries and reduced walking speed. Walking is necessary to perform many activities of daily living and is highly correlated with quality of life [1]. As such, improved walking ability is the main focus in rehabilitation for many post-stroke individuals.

Walking ability is commonly assessed by walking speed. Walking speed is dependent on propulsive force, generated during the push off phase of gate, both in healthy and stroke populations [1]. Propulsive impulse, the propulsive force integrated over time, is determined by two factors: posture of the trailing limb at push-off and ankle moment. Recent work has shown that push-off posture has a greater relative contribution than ankle moment [2].

Currently, a large body of research is focused on how to use robotic assistance to modulate the components of the propulsive impulse mechanism for rehabilitation and performance augmentation. A robotic exoskeleton that increases ankle plantarflexion torque during push off has been shown to be metabolically advantageous during treadmill and overground walking, and is currently used during robot-assisted gait training for post-stroke individuals [3], [4]. Our group is currently working on developing robotic multi-joint assistance algorithms that modulate the posture of the trailing limb at push-off to increase propulsion [5], [6]. Unfortunately, wearable exoskeletons are not always feasible for gait training of stroke survivors, due to the cost of such active devices, the burden required to wear external structures, and the technical complexity of operating such devices.

In this work, we present an alternative paradigm for goal-oriented gait training that utilizes a split belt treadmill to train both components of propulsive force generation. This is achieved by accelerating the treadmill belt of the trailing limb during the double support phase of walking. The belt acceleration introduces a fictitious inertial force that requires the ankle plantarflexors to apply greater forces during push-off to maintain their anteroposterior position on the treadmill. Moreover, assuming no modification in push-off timing, accelerations of the belt cause the trailing limb to be moved at a larger average speed that results in increases in trailing limb angle (i.e push off posture). We hypothesize that introducing subjects to this dynamic distortion condition will cause locomotor adaptation that will result in measurable after effects in the form of increased propulsive force generation when subjects are returned to a non-accelerating walking condition.

Locomotor adaptation is an error-driven, learned response to a change in environment dynamics that drives new calibrations of feedforward motor commands that persist when the environmental demands are removed [7]. Adaptation is comprised of rapid, reactive changes to environmental perturbations, and slower adaptive changes that occur over minutes to hours. Larger environmental perturbations are thought to stimulate more explicit motor control centers that drive fast reactive changes, while smaller perturbations may evoke a greater implicit response that results in longer lasting adaptated behavior [8]. Locomotor adaptation approaches have been used in gait rehabilitation to address gait asymmetry [9] and foot clearance [10], but have not been used to directly affect propulsion.

The primary purpose of this study was to test the efficacy of training propulsive force using belt accelerations in healthy individuals. We tested our paradigm at two acceleration levels (perceptible and imperceptibe) to create potentially ‘explicit’ and ‘implicit’ adaptation conditions [8]. To test the effect of our intervention on propulsive force, we collected EMG data from the calf muscles, and force plate data to measure the anterior ground reaction force (AGRF) generated during push off. To test the effect of our intervention on walking kinematics, we collected kinematic marker data to measure trailing limb angle (TLA), and evaluated subject walking speed pre and post intervention. Changes in propulsive force in healthy and stroke populations are strongly correlated to lower extremity muscle strength (increased plantar flexor muscle activity) and limb position (TLA) [2], [11]. As such, we hypothesized that there would be an increase in calf muscle activation during plantar flexion and an increase in AGRF, as well as an increase in TLA and walking speed following both intervention conditions.

## II. MATERIALS AND METHODS

A total of 19 healthy individuals participated in this study (N_*perceptible*_ = 9). The study was approved by the Institutional Review Board of the University of Delaware and each subject provided written informed consent and recieved compensations for their participation.

### A. Experimental Set-Up

Our experimental protocol is shown in Fig. 1. Participants walked for 5 minutes at a user-driven walking speed, followed by 10 minutes in the intervention condition, and a final 5 minutes at a user-driven walking speed. All user-driven walking speed conditions were conducted using a user-driven speed controller, described in Sec. II-C. Subjects walked on an instrumented dual-belt treadmill (Bertec Corp., Columbus OH, USA), while wearing four reflective spherical markers (two per greater trochanter, two per lateral malleous), and sixteen bipolar EMG electrodes (bilaterally on the tibialis anterior, lateral gastrocnemius, medial gastrocnemius, and soleous muscles). A ten camera Vicon T40-S passive motion capture system (Oxford Metrics, Oxford, UK) was used to measure marker position in space, and an OT Bioelettronica amplifier and software were used to acquire EMG. Due to system limitations, marker data was acquired only during the four periods highlighted in Fig. 1, at 100 Hz. EMG and treadmill analog force/torque data were acquired throughout the entire experimental protocol, at 10,240 Hz and 500 Hz respectively. A 24-in screen at eye level in front of the treadmill was used to provide feedback on protocol duration and as a visual target to keep subjects from looking down at the treadmill. Subjects were given noise cancelling headphones (COWIN E7 Bluetooth Headphones) that played white noise to prevent distraction from environmental sounds.

**Fig. 1.**
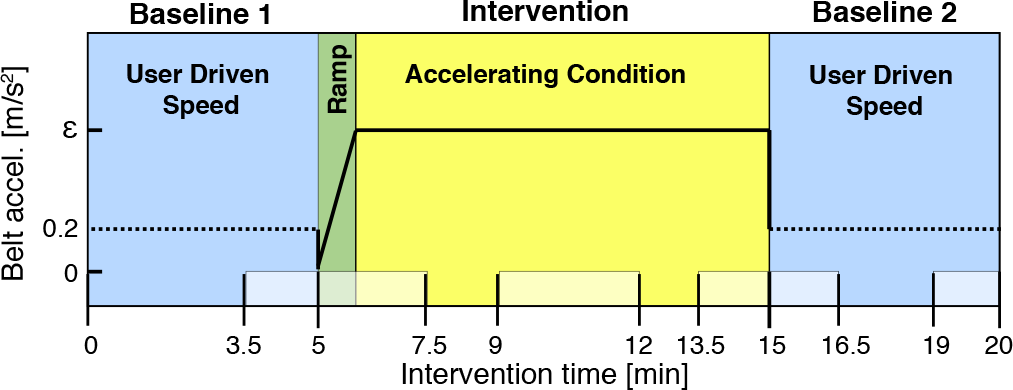
Experimental Protocol Schematic. Belt accelerations are shown on the y-axis, where *∊* signifies the magnitude of accelerations applied during intervention (either 2 m/s^2^ or 7 m/s^2^). A minute long “ramp” phase (green), in which accelerations linearly increased in magnitude from 0 to *∊*, was used to gradually introduce the intevention condition. For the UDTC, maximum belt accelerations were capped at 0.2m/s^2^. Highlighted phases at the bottom signify periods in which kinematic marker data were collected.

### B. Self-selected walking speed

A preliminary set of trials were conducted to determine subjects self-selected walking speed (SS-WS) prior to intervention. Participants walked on the treadmill at an initial speed of 0.5 m/s that was gradually increased in intervals of 0.03 m/s by the experimenter until the participant verbally indicated the treadmill had reached their SS-WS. The same procedure was repeated with the treadmill beginning at 1.8 m/s and decreasing in increments of 0.03 m/s until the subject indicated their SS-WS had been reached. Each procedure was repeated three times and subjects SS-WS was calculated as the average between the six measured speed values [6].

### C. User-Driven Speed Controller

In standard treadmill subject walking, walking speed is restricted to the constant velocity imposed by the treadmill. In our intervention that seeks to modify subjects walking speed, a treadmill that operates at a constant velocity–and thus restricts changes in walking speed–is impractical, and likely to eliminate the after effects any dynamic distortion may induce. To address this, we employed a user-driven treadmill controller (UDTC) that changes the velocity of the treadmill belts in response to changes in the subjects walking behavior [12].

The UDTC changes speed based on an empirically weighted combination of the following three gait parameters: change in AGRF, step length, and position relative to the center of the treadmill. For example, if subjects produced greater AGRF, increased their step length, or walked further forward on the treadmill, the treadmill speed would increase. Conversely, decreases in AGRF, step length, or movement to the back of the treadmill would decrease speed. The maximum belt acceleration was set to 0.2 m/*s*^2^, and was previously tested to ensure subject safety and comfort [12]. Prior to our experimental protocol, participants were trained on the UDTC condition. Participants were started at their SS-WS and given up to 10 minutes to familiarize themselves with the UDTC. In line with previous results, several subjects increased their walking speed on the UDTC from their initial SS-WS. Consequently, for our experimental protocol, the initial baseline period of walking was set to at least 5 minutes, but was continued untill subjects held a constant velocity (± 0.05 m/s) for one minute. The Baseline 1 (Fig. 1) condition of our protocol was then defined as the period spanning the one minute of steady state walking, and the previous four minutes prior to acheiving a steady state gait speed.

### D. Accelerating Controller

To train increases in propulsive force, we developed a controller capable of accelerating the treadmill belt of the trailing limb during the double support phase of gait and returning it to a nominal speed during the swing phase of gait (Fig. 2). The rationale behind the selection of this dynamic distortion (acceleration), was to attenuate subjects propulsive force by introducing a fictitious inertial force in the opposite direction. Moreover, assuming that subjects do not modify their push-off timing, the foot on the accelerating belt will move at a larger average speed, causing TLA to increase. In this way, we hoped to target the both gait mechanisms (ankle moment and posture of the trailing limb at push-off) that control propulsive force generation.

**Fig. 2.**
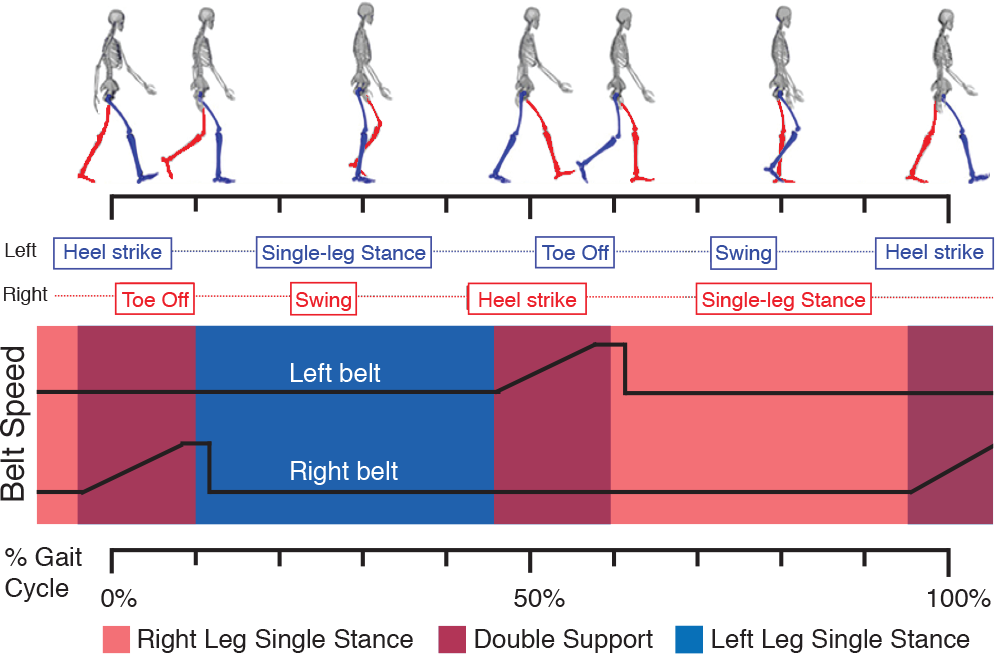
Belt accceleration during intervention as a function of gait cycle

Push off occurs at the end of the double support phase of gait that typically lasts for 100-150 ms. While double support can be detected in real time using force-plate data, delays in the measurement and actuation components of our treadmill system that exceeded 150 ms between detection of dual support and the execution of an acceleration command made using real time detection impossible. Instead, we developed a simple algorithm to predict when push off would occur based on the prior gait cycle, that included an anticipation factor to account for our system delays. This controller commands the belt to accelerate at a time *t*, measured relative to the instant of heel strike of the leg to be accelerated, defined by

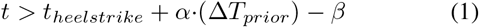

where *t* is the current time, *t_heelstrike_* is the heelstrike time of the leg currently in stance, Δ*T_prior_* is the time between the previous left and right heel strikes that serves as a prediction for the time untill double support will occur, and *β* is the anticipation factor. Once the amount of time elapsed since heel strike exceeds *α·*(Δ*T_prior_*) *− β* the acceleration signal is sent. A weighting of *α* = 1.175 and *β* = 0.185 s were determined empirically on a seperate group of 10 individuals to confirm accelerations occured during pushoff at a reasonable range of speeds (0.7 - 1.4 m/s). Accelerations were saturated by a speed increase limit of 0.6 m/s to ensure subjects safety. 100 ms after detection of toe-off, the controller decelerates the belt back to the nominal speed during the swing phase (Fig. 2).

#### 1) Data Preprocessing

EMG, kinematic marker data, and forceplate data were acquired on 3 separate systems and time synced via a common signal. VICON marker position data were fed into a standard Visual3D pre-processing pipeline, which included i) manual labelling of markers; ii) interpolation of missing marker data with a third order polynomial fit for a maximum gap size of five samples; and iii) low-pass filtering at 6 Hz with a 4th order zero-shift Butterworth filter [6]. Kinematic marker data for one subject in the Imperceptible group was excluded due to excessive marker drop out. Force-plate data recorded in Matlab were filtered with a 4th order zero-shift low-pass Butterworth filter at 200 Hz [6]. EMG data were bandpass filtered at 10-500 Hz, rectified, and the envelope was taken using a lowpass zero-shift 4th order Butterworth filter with a cut off frequency of 50 Hz [13].

#### 2) Data Analysis

Force-plate data were used to segment EMG data by gait cycle. Gait cycles were defined by heel strike events, determined as the instants at which the vertical ground reaction force exceeded 25 N and remained above 25 N for at least 200 ms. AGRF was calculated from the force-plate data as the area under the positive (anterior) portion of the curve of the ground reaction force (Fig. 3) [1]. TLA was defined as the angle between the straight line connecting the greater trochanter and the lateral malleous of the trailing limb and the vertical axis of the lab at the time of peak AGRF (Fig. 3) [1]. Gait speed (GS), measured continously by the treadmill, was sampled at right and left heelstrike events to get a measure of speed per step with a consistent sampling rate as used in AGRF and TLA measures.

Because we expected our two intervention groups to elicit adaptative behaviors with different timescales, we assessed the progression of after effects following intervention. To this end, the second baseline period (BL2) was partitioned into bins of 20 strides, and the distribution of gait parameters within each bin was used for subsequent statistical analysis.

**Fig. 3.**
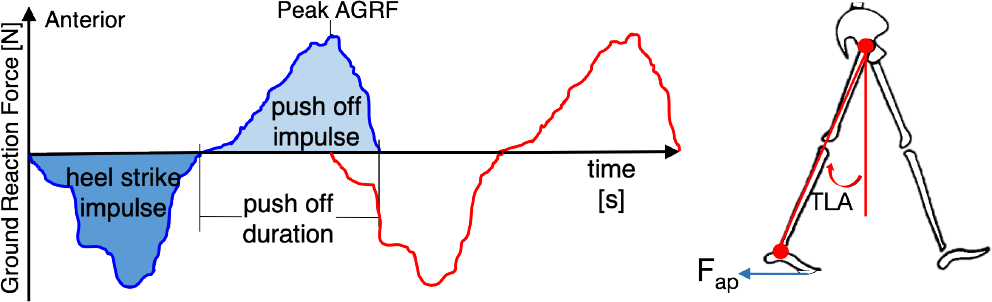
Definition of gait parameters AGRF (left) and TLA (right)

To prepare EMG data for statistical analysis, EMG data of the tibialis anterior, lateral gastrocnemius, medial gastrocnemius, and soleus muscles were segmented and linearly resampled to [0 - 100] percent of gait cycle. Technical difficulties associated with signal synchronization between force plate and EMG acquisition, as well as noise in our EMG acquisition system affected data in 5/9 subjects in the Perceptible intervention group and 1/10 subjects in the Imperceptible group. Consequently, only the Imperceptible intervention group had sufficient data for group analysis.

### E. Statistical Analysis

#### 1) Group Analysis

To evaluate the effects of intervention, and intervention type (Perceptible vs. Imperceptible) on subject gait parameters, we used three 2-way mixed effects ANOVAs, one per each outcome measure (AGRF, GS, and TLA). For AGRF and GS, the within-subjects factors (experimental condition) included baseline 1 (BL1), defined as the mean of last minute of baseline 1 walking, and BL2 1-9, defined as the mean values measured during each of the nine contiguous 20-stride bins that spanned strides 1-180 in BL2. Due to our experimental limitations in data collection, for TLA, we defined BL1 as before, but used only five 20-stride bins that corresponded to strides 1-60 (bins 1-3) and the last forty strides (bins 4-5) of BL2. Bins in early BL2 were included to quantify the immediate effects of the intervention, while bins in late BL2 were used to evaluate the sustained effects of the intervention. For each outcome measure, the mean was obtained by averaging right and left leg measures. Post-hoc testing (paired-t-tests) was used to quantify group level effects pre and post-intervention, by comparison of the gait parameters measured in BL1 to each BL2 bin.

As EMG data are measured as 1D timeseries, we used the software SPM1d to conduct a one-way repeated measures ANOVA to test for significant effect of the intervention on muscle activity at any instant over the gait cycle [14]. We included 5 experimental factors in our ANOVA: BL1, Ramp, Intv, Early BL2, and Late BL2. Because of the intrinsic noise characteristics in EMG recordings, we used a simplified binning strategy to define the within-subject factors as follows: BL1, defined as the last minute of baseline 1 walking; Ramp, defined as the first 20 strides of intervention; Intv, defined as the last 20 strides of intervention; Early BL2, defined as the first 20 strides following intervention; and Late BL2, defined as the final 20 strides in the baseline 2 condition. Post-hoc paired t-tests were used to evaluate the effect of each within subject factor on EMG activations. It is important to note that EMG measurements were the only gait parameter that directly measured subjects response to dynamic distortions that could be accurately measured during intervention. In fact, during intervention GS is imposed, AGRF measurements are inaccurate due to the non-inertial conditions imposed by belt accelerations, and peak AGRF is necessary to determine our TLA measure [1], [15].

#### 2) Within Subject Analysis

The ANOVA analyses described above were adapted for evaluation of effects of the intervention at the individual subject level. For gait parameters (AGRF, GS, TLA) a 2-way mixed effects ANOVA was used, with within-subject factors defined in the same way as the group level analysis (with the exception of within-bin averaging). To account for gait asymmetries, single-subject ANOVAs included a second factor: leg (right or left). Post-hoc testing via two-sample t-test was used to quantify the effects of intervention by comparison of the gait parameters measured in BL1 to each BL2 bin. To characterize the variability in response to intervention, the ANOVA results were used to classify subjects response to intervention. Subjects were classified positive responders if they had a significant increase in BL2 compared to BL1 in the first half of BL2 bins (most directly associated with after-effects of intervention), and non - responders if they had significant decreases or no change in walking behavior.

The same ANOVA used for group-level EMG analysis was run at the single-subject level for all viable subjects in both intervention groups. Post-hoc two sample t-tests were used to determine the effect of each intervention condition (BL1, Ramp, Intv, Early BL2 and Late BL2) on EMG activations.

## III. RESULTS

### A. Group Analysis

#### 1) Propulsive Force Generation

##### Anterior Ground Reaction Force

The ANOVA fit the data with an 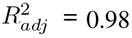. A significant effect of intervention was detected (*p* < 0.001), however the interaction between intervention type and the within-subject factor failed to achieve our selected significance threshold (*p* = 0.10). Post-hoc testing revealed a significant increases in AGRF (*μ* = 462 Nm*·*s) in all BL2 bins compared to BL1 across intervention groups (*p* < 0.05). For the Perceptible group, the mean increase in AGRF measured across all BL2 bins was 631 Nm*·*s, and was larger than the mean increase measured in the Imperceptible group (294 Nm*·*s). AGRF measured in later bins (8-9) was significantly greater than in early bins (1-3) across intervention groups (200 Nm*·*s). For the Perceptible intervention group, all BL2 bins were significantly greater than BL1, and late BL2 bins 8-9 were greater than early BL2 bins (1-2) (Fig. 4, top). For the Imperceptible group, all bins except 2 and 5 showed significant increases in AGRF.

**Fig. 4.**
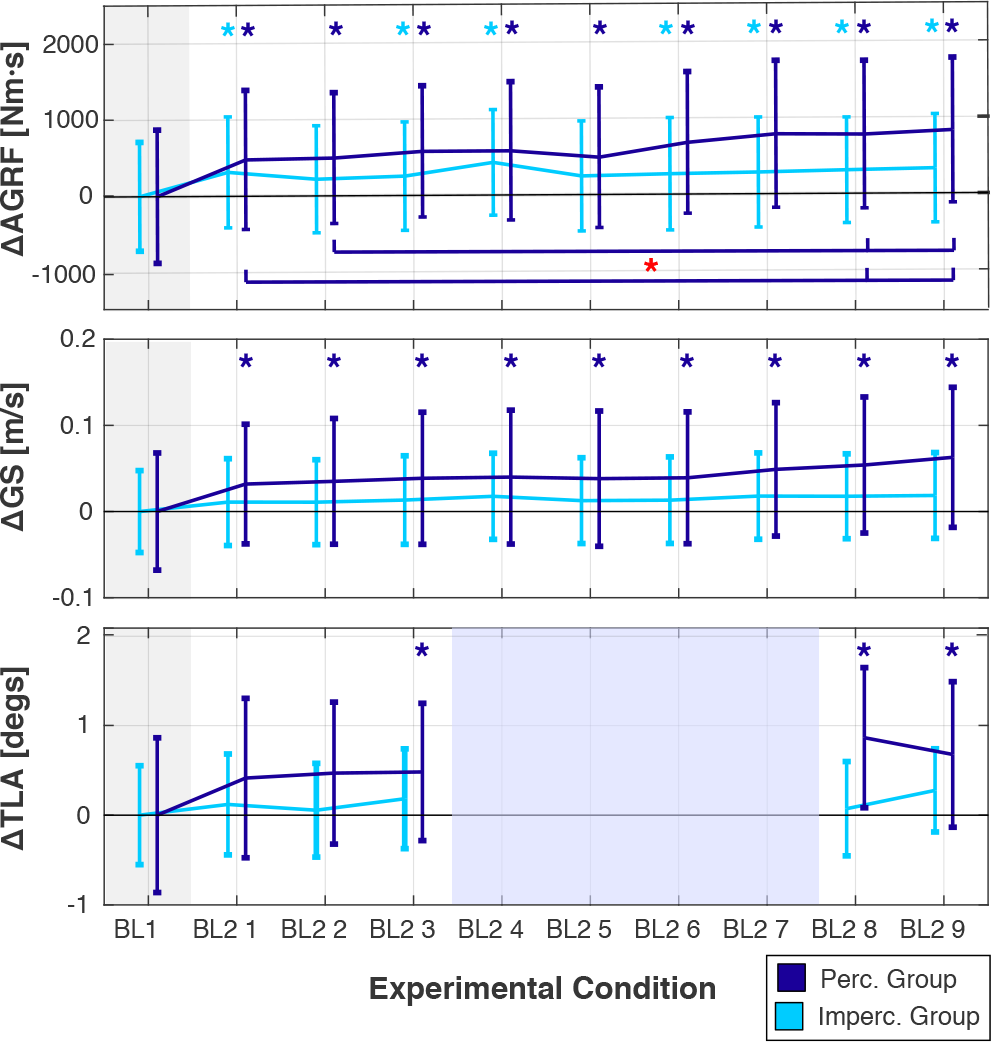
Group level pre and post intervention change in AGRF, GS, and TLA. Each bar plot displays the mean and standard error in each bin. Astrics at the top of each plot display significant change between BL2 bins and BL1. Astrics at the bottom display significant change between denoted BL2 bins.

##### EMG

The repeated measures ANOVA in SPM1d of the Imperceptible intervention group showed a significant effect of intervention on plantar flexion muscles activation centered around 45-55% of gait cycle, that is roughly aligned with the double support phase of gait (Fig. 7). Post-hoc analysis revealed this effect was driven by increases in muscle activation during intervention that were sustained at a smaller amplitude in baseline 2 walking.

#### 2) Walking Kinematics

##### Gait Speed

The ANOVA fit the data with an 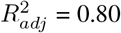. A significant effect of intervention was determined (*p* = 0.017), but the interaction between intervention type and the within-subject factor failed to achieve our selected significance threshold (*p* = 0.30). Post-hoc testing revealed a significant increase in GS (*μ* = 0.03 m/s) in all BL2 bins compared to BL1 across intervention groups (*p* < 0.05). Specifically, for the Perceptible group, the mean increase in GS measured in all BL2 bins was equal to 0.043 m/s, and was larger than the mean increase measured in the Imperceptible group (0.015 m/s). For the Perceptible intervention group, GS in all BL2 bins was significantly greater than in BL1 (Fig. 4, middle). For the Imperceptible group, no significant change in GS was detected.

##### Trailing Limb Angle

The ANOVA fit the data with an 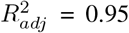. The effect of intervention on TLA failed to achieve our selected significance threshold (p = 0.06). Mean TLA increase in BL2 bins compared to BL1 across intervention groups was 0.30 deg, 0.48 deg for the Perceptible group, and 0.18 deg for the Imperceptible group. Post-hoc testing revealed a significant increase in TLA (*μ* = 0.47 deg) in late BL2 bins (4 and 5) compared to BL1 across intervention groups (*p* < 0.05). In the Perceptible intervention group, Late BL2 bins 3-5 were significantly greater than BL1 (*p* < 0.05) (Fig. 4, bottom). For the Imperceptible group, no significant change in TLA was detected.

### B. Within Subject Analysis

#### 1) Propulsive Force Generation

##### Anterior Ground Reaction Force

The ANOVA rejected the null hypothesis of equal means between conditions for all subjects. Seven subjects in the Perceptible group and eight the Imperceptible group were classified as positive responders that had significant increases in AGRF compared to BL1 Two subjects in each group had significant decreases in AGRF and were classified as non-responders.

##### EMG

Subject level ANOVA of EMG data showed significant effects of intervention on muscle activation for 12/13 subjects tested in at least one muscle group. Subjects with the largest effects in EMG activation corresponded to individuals with the largest changes in AGRF: the two individuals with the largest increases in AGRF in each group had the largest positive increases in EMG activations during and following intervention. Similarly, the two individuals who had significant decreases in AGRF following intervention had decreased EMG activations during intervention, that remained decreased in BL2 walking. Fig. 6 shows the two sample t-test results for EMG recorded in the lateral gastrocnemius muscle for representative positive and negative non-responders in both intervention groups.

**Fig. 5.**
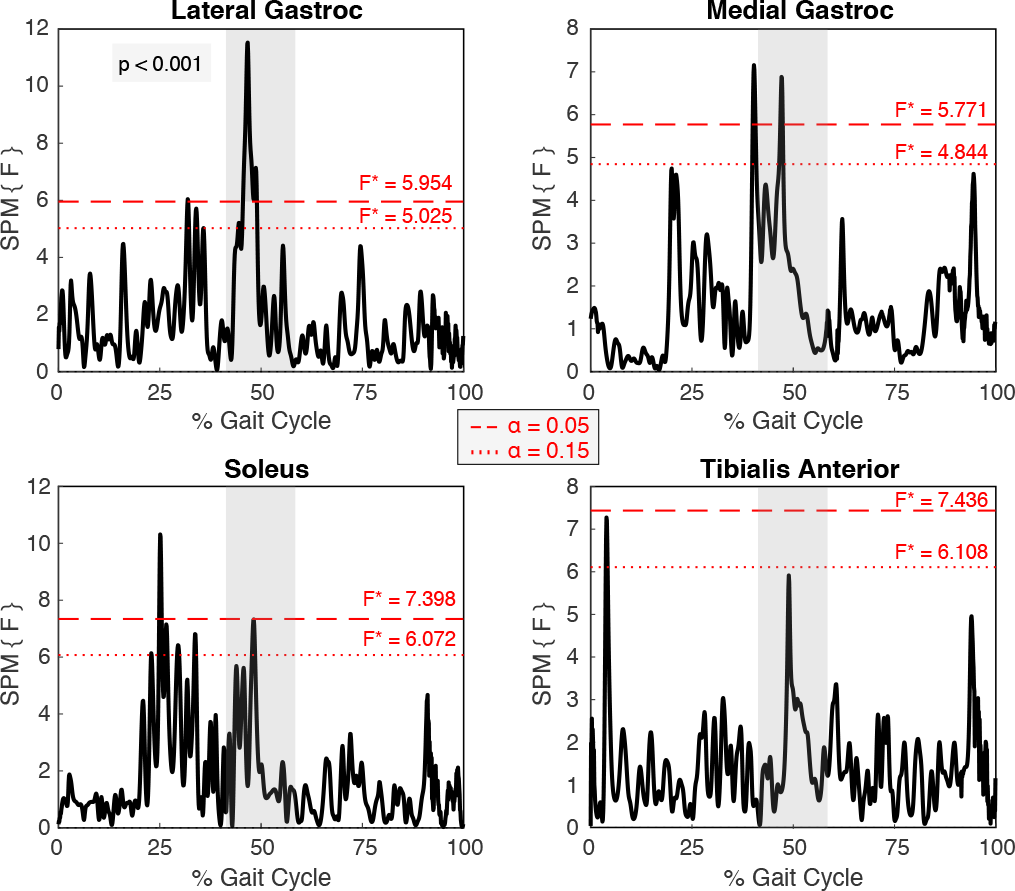
Group ANOVA results of the Imperceptible intervention on EMG as a function of gait cycle. There was a significant effect of intervention on calf muscle activity. Post-hoc analysis showed the effects centered around 45-55% of gait cycle (double support phase of gait) were due to increases in muscle activation during intervention that were sustained in BL 2 walking.

**Fig. 6.**
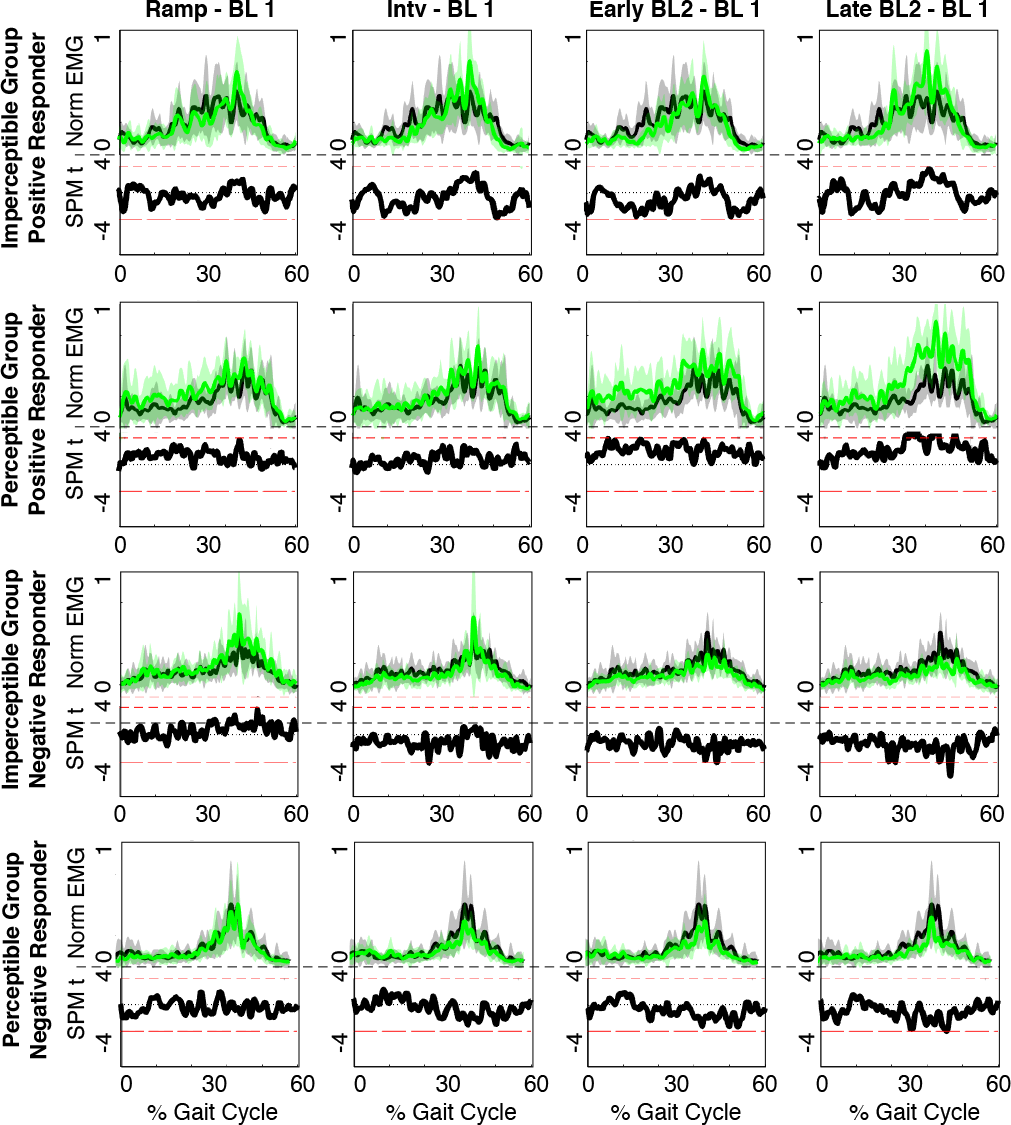
Within subject analysis of responders Lateral Gastrocnemius EMG activity. The top portion of each split plot displays the mean and standard deviation (shaded area) of the EMG tracks (Grey is BL1, Green is experimental condition). The bottom portion reports T-scores (black) of the post-hoc two sample t-tests comparing each experimental condition to BL1. Dashed red lines display the *α* =0.05% boundaries.

#### 2) Walking Kinematics

##### Gait Speed

The ANOVA rejected the null hypothesis of equal means between conditions for all subjects. Seven subjects in the Perceptible group and seven in the Imperceptible group were classified as positive responders that had significant increases in GS compared to BL1. Two subjects in the Perceptible group, and three subjects in the Imperceptible group had significant decreases in GS and were classified as non-responders. One subject in the Perceptible, and two subjects in the Imperceptible group who were ‘positive responders’ based on initial change had significant decreases in the later half of BL2 bins considered.

##### Trailing Limb Angle

The ANOVA rejected the null hypothesis of equal means between conditions for all subjects. Seven subjects in the Perceptible group and five in the Imperceptible group were classified as positive responders that had significant increases in TLA. Two subjects in the Perceptible, and four subjects in the Imperceptible group were calssified as non-responders that had significant decreases in TLA.

## IV. DISCUSSION & CONCLUSION

We presented a novel paradigm used to train two components of propulsive force generation during walking (push-off posture and ankle plantarflexor moment), based on the application of belt accelerations to the trailing limb during the double support phase of gait. In our first human-subject experiment evaluating the effects of this intervention, we exposed two groups of subjects to belt accelerations at two different magnitudes (Perceptible and Imperceptible).

Our analysis showed that, at the group level, our intervention elicited increases in propulsive force generation but not in push-off posture. Statistical analysis of our propulsive force generation measures showed that muscle activity increased both during and following intervention, and that AGRF increased following intervention. The most consistent effect of our intervention across both intervention groups was an increase in propulsive force, quantified by change in AGRF. Increases in AGRF translated to modest changes in walking kinematics that included increases in WS and TLA. From these results, it appears that our intervention acts predominantly on the ankle moment mechanism of propulsive force generation, and to a lesser extent on TLA.

The Perceptible group had larger initial increases in AGRF compared to the Imperceptible group, consistent with expected behavior following explicit adaptation. Non-consistent with explicit adaptation was the sustained increase in AGRF across the BL2 walking period, that remained greater than the Imperceptible group at all time points tested. The Imperceptible group also had larger variability in subject response to intervention compared to the Perceptible group. These results taken together suggest that the effects of our experimental protocol may not be dependent on the adaptation processes engaged. Instead, it is possible both protocols stimulate implicit adaptation processes in a dose dependent manner, and that the Imperceptible group accelerations were insufficient to cause reliable after effects.

Overall, our paradigm significantly increased propulsive force generation in 78% of subjects in the Perceptible group and 80% in the Imperceptible group, that translated to increases in gait speed in 66% and 50% of subjects respectively. Our results indicate the Perceptible intervention is a better candidate for continued research in training propulsion, as subjects in this group had more consistent after-effects, larger increases in all gait parameters tested, and sustained change in walking behavior following intervention.

## V. ACKNOWLEDGMENTS

We acknowledge support from the University of Delaware Research Foundation grant no. 16A01402, and from startup funds by the University of Delaware. We thank Andrew Borowski for his collaboration on the accelerating controller, and Nicole Ray for her collaboration on the UDTC.

